# A Semi-Automated MEA Spike sorting (SAMS) method for high throughput assessment of cultured neurons

**DOI:** 10.1101/2025.02.08.637245

**Authors:** Xiaoxuan Ren, Carissa L. Sirois, Raymond Doudlah, Natasha M Mendez-Albelo, Aviad Hai, Ari Rosenberg, Xinyu Zhao

## Abstract

Neurons derived from human pluripotent stem cells (hPSCs) are valuable models for studying brain development and developing therapies for brain disorders. Evaluating human-derived neurons requires assessing their electrical activity, which can be achieved using multi-electrode arrays (MEAs) for extracellular recordings. Because each electrode channel generally detects activity from multiple neurons, resolving the activity of single neurons requires a process called spike sorting. However, currently available spike sorting methods are not optimized for the analysis of hPSC-derived neurons, and require complex workflows and time-consuming manual intervention. Here, we introduce a **S**emi-**A**utomated **M**EA **S**pike sorting software (SAMS) designed specifically for low-density MEA recordings of cultured neurons. SAMS outperforms commercially available automated spike sorting algorithms in terms of accuracy and greatly reduces computational and human processing time. By providing an accessible, efficient, and integrated platform for spike sorting, SAMS enhances the resolution and utility of MEA in disease modeling and drug development using human-derived neurons.

**Highlights:**

- SAMS is designed and optimized for high throughput analysis of hPSC-derived neurons.
- SAMS is more efficient and accurate compared to recommended spike-sorting software.
- SAMS resolves phenotypic differences previously not observed without spike sorting.
- SAMS is an open-source software.

## INTRODUCTION

Human pluripotent stem cells (hPSCs), including embryonic (hESCs) and induced pluripotent stem cells (hiPSCs), have provided useful experimental models for defining human-specific biology and disease mechanisms, especially for brain disorders.^1–4^ To evaluate the development and function of hPSC-derived neurons, it is important to assess their electrophysiological properties, particularly their spiking characteristics. Multi-electrode arrays (MEAs) are a widely used tool for simultaneously recording from many neurons *in vivo,*^5–8^ *ex vivo,*^9^ or *in vitro.*^10–13^ MEAs detect extracellular field potentials generated by cells proximal to the electrodes within the tissue (*in vivo*), or near to the tissue or cells plated onto the electrodes (*ex vivo* and *in vitro*, respectively). Recently, MEAs have been used to assess both the intrinsic properties and network activities of hPSC-derived neurons, allowing for relatively high throughput assessment of disease genes and drug-like compounds.^14–17^ However, widely used low-density MEA recordings often capture the combined activity of multiple neurons on an individual electrode. Therefore, to accurately characterize neuronal properties of hPSC-derived neurons, it is crucial that the activity of individual neurons are isolated. This is done by sorting spikes into groups based on similar spike waveform characteristics.

Currently, many spike sorting pipelines exist for both high-density and lower-density MEAs (**Table S1**). However, the majority of these pipelines have been created to analyze either *in vivo* recordings or primary neuronal cultures from non-human species,^18^ and have not been optimized for hPSC-derived neurons that have different physiological properties, affected by both differentiation methods and disease states.^19–23^ Furthermore, these pipelines require coding skills to either run the spike sorting algorithms or implement bulk processing over many datasets, and some pipelines require the use of additional software tools to perform analyses on the spike-sorted data. These pipelines thus require time consuming manual checking and correction of the sorted spikes on each electrode,^24–29^ which limits their application to the analysis of hPSC-derived neurons in 48 or 96 well MEA plates which contain hundreds of electrodes on a single plate.

To overcome the challenges of current spike sorting approaches, we developed a **<u>S</u>**emi-**<u>A</u>**utomated **<u>M</u>**EA **<u>S</u>**pike sorting software (SAMS) for low-density MEA data collected from hPSC-derived neurons using the Axion MEA platform. Our goal was to create an open-source end-to-end spike sorting and analysis tool that was user friendly and accessible to more researchers, especially those without computational expertise. SAMS is user interface based and requires no coding to implement spike sorting routines or to change parameters. The workflow is easy to follow, performing automated spike sorting, and characterizing the spiking parameters of individual neurons. Importantly, SAMS flags electrodes that may have been incorrectly sorted, prioritizing which electrodes that needed to be manually verified, reducing the time required to validate the sorted spikes. We compared the spike sorting performance of SAMS with Plexon’s Offline Sorter (OFS) software, which is the current software recommended for analyzing low-density MEA recordings obtained using the Axion platform. We found that SAMS not only outperformed OFS on spike sorting accuracy but also decreased the amount of manual validation time. We then used SAMS to reanalyze two previously published MEA datasets of hPSC-derived neurons where no spike sorting had been performed.^30,31^ The SAMS pipeline not only revealed similar differences between experimental conditions as previously published but also uncovered additional differences when examining neurons grouped by their levels of bursting activity. Therefore, SAMS enhances the resolution and utility of MEA recordings in disease modeling and drug development using human-derived neurons.

## RESULTS

### Overview of spike sorting recommendations for the Axion MEA platform

Currently, the suggested pipeline for spike sorting and analysis of electrophysiological recordings obtained using Axion Biosciences’ MEA system involves multiple complex steps performed across several software tools (**Figure 1A**). First, Axion’s Axis Navigator software is used to generate a proprietary spike file from the raw recording data file. The spike file contains the spike times and voltage waveforms measured for each electrode with the waveforms aligned at their peaks to facilitate spike sorting. Next, the spike file is converted to the Neuroexplorer file format in the Axion Data Export Tool, which preserves important metadata such as the plate format and allows the file to be analyzed by other commercial software. Command scripts must then be written to batch process the spike sorting of multiple files in OFS, or users can individually process single files in OFS using manual, semi-automated, or automated spike-sorting algorithms. This is then followed by manual inspection and sorting of all electrode channels to identify and assign individual waveforms to single neurons. This is a time-consuming process that depends on the total number of waveforms (reflecting the amount of activity recorded by the electrodes) that need to be sorted, the number of putative units identified, and how distinct the waveform characteristics are that define each unit. Finally, the sorted spike outputs are exported and then analyzed using Axion’s Neural Metric Tool, which allows the user to obtain per-unit and per-well spike and bursting data, as well as assess functional connectivity and synchrony. This workflow burdens users by necessitating coding proficiency for scripting, while also relying on time-intensive manual validation across channels. Alternatively, users can write their own custom pipelines or employ previously published pipelines (**Table S1**) to perform spike sorting using MATLAB or Python.

**Figure 1.**
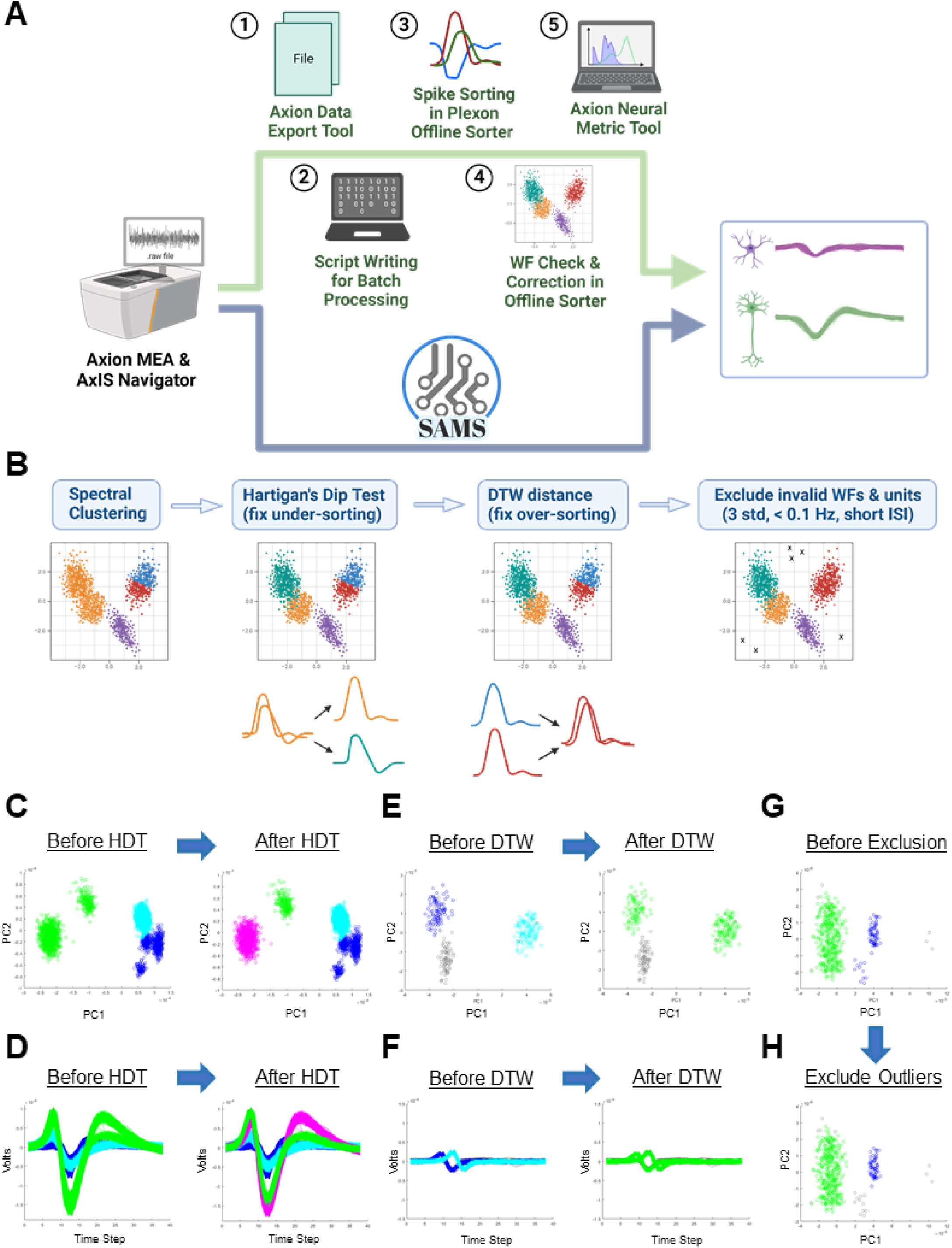
Comparison of standard spike sorting workflow and SAMS. A. Top: The standard pipeline for spike sorting and analysis of recordings from Axion’s MEA system. (1) MEA data is converted from the proprietary spike file format to a file format compatible with Plexon’s Offline Sorter (OFS) software using Axion’s Data Export Tool. (2) Users manually write scripts for batch processing of files in OFS. (3) Automatic spike sorting is performed in OFS. (4) Manual inspection and correction of spike sorting results for each electrode in OFS. (5) The sorted spike outputs are analyzed using Neural Metric Tool to assess functional connectivity. The workflow produces individual unit analysis and network activity visualizations. **Bottom:** Same pipeline using SAMS. SAMS streamlines the processes shown above by directly interfacing with Axis Navigator output files and performing automatic batch processing through four steps detailed in panel B. **B.** Schematic illustrating the four stages of MEA data processing performed by SAMS: spectral clustering to perform initial waveform clustering, Hartigan’s Dip Test (HDT) to fix under-sorting, DTW distance calculation to fix over-sorting, and exclusion of invalid waveforms based on deviation from the average waveform shape, spike frequency criteria, and short inter-spike interval (ISI). **C-H.** PCA plots and waveform traces from analyzed data showing example electrodes corrected by different stages of the SAMS pipeline. **C,D.** Correction of an under-sorted electrode using Hartigan’s Dip Test. **E,F.** Correction of an over-sorted electrode using DTW distance. **G,H.** Removal of outlier waveforms.

However, these approaches still require coding proficiency for scripting and, importantly, are not intended or optimized for use with low density recordings from hPSC-derived neurons.

### Overview of spike sorting using SAMS

To address the inefficiencies described above, we developed SAMS as an integrated one-stop platform enabling automated batch processing of MEA datafiles from Axion directly into analyzed output files (**Figure 1A**). SAMS provides an intuitive graphical user-interface to eliminate the need for scripting, while integrating manual spike sorting tools. By consolidating multiple manual steps that are required for other pipelines, SAMS offers a streamlined workflow to accelerate data processing and analysis. The simplified user experience lowers adoption barriers, enabling bench scientists without specialized computational skills to readily incorporate high fidelity information on neuronal encodings and circuits into their research.

SAMS employs a multi-step spike sorting pipeline for robust classification (**Figure 1B-H**). First, spectral clustering^32^ is used to perform initial unit assignments based on detected spike waveform features. This unsupervised step maximizes between-cluster variance while minimizing within-cluster variance. Second, Hartigan’s Dip Test (HDT)^33^ is used to assess unit purity by sampling and comparing the empirical cumulative distribution of the nearest neighbors for each cluster. Clusters failing the dip test are recursively split to refine sorting (**Figure 1C,D**). Third, dynamic time warping (DTW)^34^ is used to resolve over-sorting errors by merging units with temporally warped matching spike waveforms (**Figure 1E,F**), which can occur when spike detection is triggered using a simple threshold crossing, which is defined by the user (default threshold: 1.2; **Figure S1**). Finally, a series of sequential steps is used to identify and remove the following three types of invalid waveforms: (1) waveforms exceeding three standard deviations from each unit’s mean template (i.e., averaged waveform), (2) waveforms from low firing rate units (default threshold: < 0.1 Hz)^23^ (**Figure 1G,H**), and (3) waveforms with inter-spike intervals (ISIs) falling within the user-defined refractory period (default: 1.5 ms). By integrating quality control metrics at each step – testing cluster integrity, merging probable over-splits, and removing outliers or false positives – SAMS balances the demands of spike sorting accuracy and manual intervention. The multi-tier workflow maximizes the yield of high confidence isolated single-units essential for revealing biological insights and allows the user to customize parameters at each stage of the pipeline to optimize sorting of their data (**Table S2**). SAMS also returns an output file containing the parameter settings used for each analysis to facilitate transparent reporting of methods and allow users to verify that settings were consistent between analysis batches.

### SAMS provides a user-friendly interface containing key information of output data

To enable users to customize their analyses, SAMS includes panels (**Figure 2A**) for tuning single unit burst detection parameters at the single-electrode level (minimum spike count; maximum intra-burst inter-spike interval (ISI)) as well as network burst detection at the multi-electrode level (spike counts; ISIs; and percentage of electrodes participating). While the default parameters are set to be consistent with the defaults applied by the Axion system and used by most researchers, the capacity to modify these parameters provides researchers additional flexibility in analyzing their data (**Figure 2A; Table S2**).

**Figure 2.**
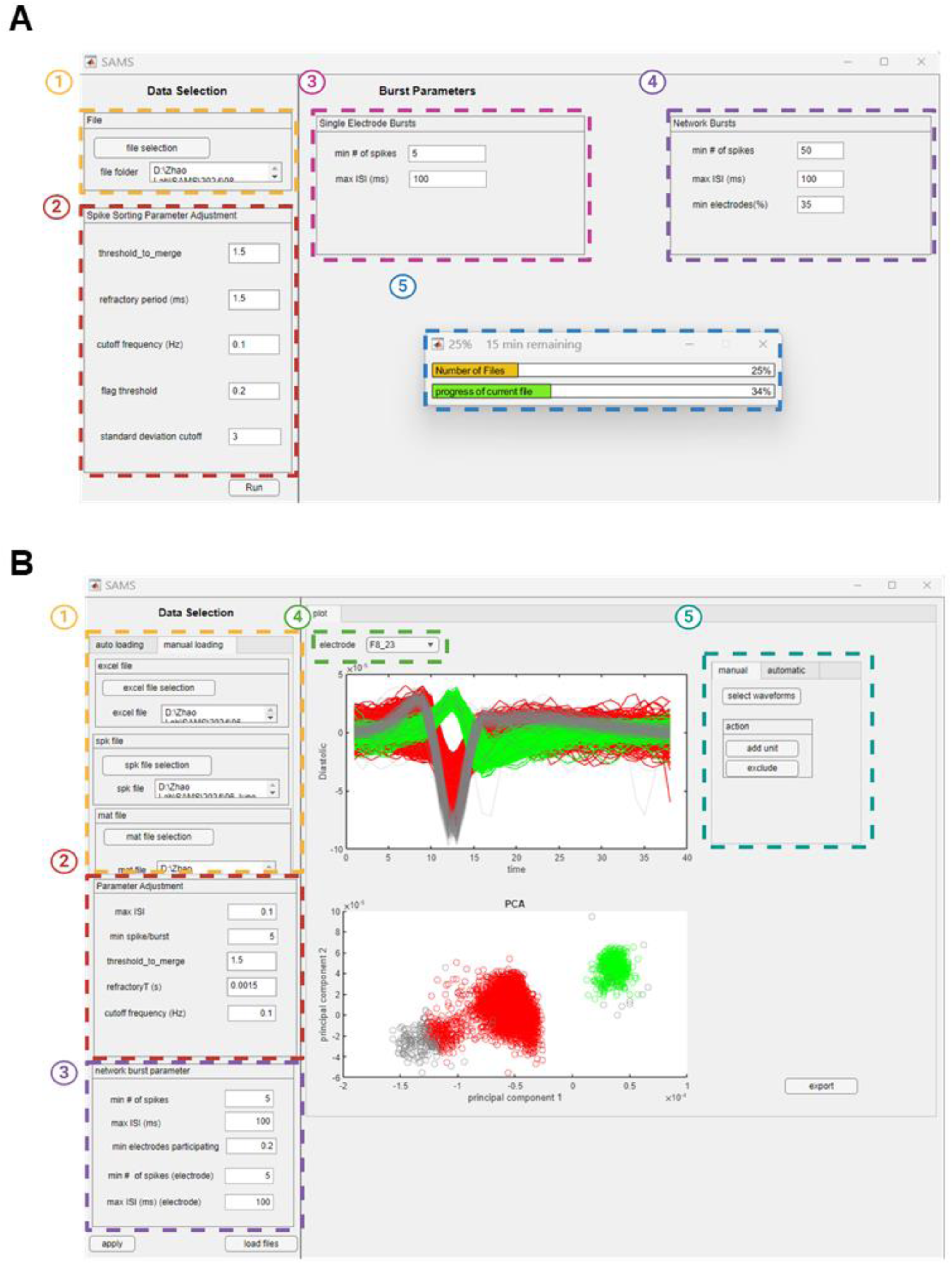
SAMS Graphical User Interfaces and Features. **A.** SAMS graphical user interface for automatic batch spike sorting. The SAMS interface facilitates efficient batch processing of spike sorting data. Key components include: (1) File Selection: User specifies the directory containing files for analysis. (2) Spike Sorting Parameter Adjustment: Parameters related to the spike sorting process can be modified by the user (**see Table S2**). (3) Single Electrode Burst Parameters: Configurable minimum spike count and maximum ISI for individual electrodes. (4) Electrode-level Network Burst Parameters: Settings for minimum spike count, maximum ISI, and minimum percentage of participating electrodes in network bursts. (5) Progress Indicator: Real-time feedback on file processing status and individual electrode analysis within the current file. This interface streamlines the spike sorting workflow, allowing users to efficiently process multiple files with tailored parameters. See Table 2 for more information. **B.** SAMS graphical user interface for Manual Correction of Spike Sorting Results. This interface facilitates precise refinement of automatic spike sorting outputs. Key components include: (1) Data Selection: Users can load previously processed files, including Excel files with unit analysis and network burst data, spike files used in spike sorting, and Matlab SAMS output files containing processed sorting information. (2,3) Parameter Adjustment: Allows users to align manual correction parameters with those used in automatic sorting, ensuring consistency. (4) Electrode Selection: A dropdown menu enables users to choose specific electrodes for manual correction. The interface displays both waveform plots and corresponding principal component analysis (PCA) visualizations for the selected electrode, enabling informed decision-making during the correction process. (5) Correction Tools: Options for manual or automatic adjustments, including the ability to select waveforms and add units or exclude data. Note: Data shown are not the final sorted waveforms for the example electrode.

While designed for primarily automated batch processing, SAMS also enables user-friendly manual verification and refinement by providing a multi-faceted graphic user interface (**Figure 2B**) and output files (**Figure 3A**) for efficient review of sorting quality, detailed single-unit analysis, and assessment of network activity patterns. Initial automated sorting outputs are compiled into a convenient slide deck results file (**Figure 3B**), which visually summarizes unit assignments across all electrode channels for inspection. Results from each electrode are shown on individual slides, which include waveform plots of the identified units, a principal component analysis (PCA) plot, and the number of valid and invalid waveforms. Detailed results for each identified unit are contained in a spreadsheet which contains a summary of the parameters used in the analysis (**Figure 3C**), electrode-level network activity metrics, single unit metrics such as mean firing rate and bursting activity metrics, and an electrode statistics tab which provides summary-level data for each electrode such as total number of spikes. SAMS also returns two lists of flagged electrodes that have been determined by the algorithm as possibly being incorrectly sorted: the Possible Multi-Unit list and the Over-Excluded Unit list (**Figure 3D**). Electrodes are added to the Possible Multi-Unit list if they fail HDT or if they are below the DTW Distance threshold provided by the user, and are added to the Over-Excluded Unit list if the number of waveforms excluded from a given electrode exceeds the threshold (percentage of total waveforms) specified by the user. If channels show suboptimal separation, the user can load files back into SAMS’ manual sorting interface which supplies spike waveforms and PCA feature plots (**Figure 2B**). Using an intuitive click-and-drag selection tool with “add unit” and “exclude” buttons, users can assign selected spike waveforms to new or existing units.

**Figure 3.**
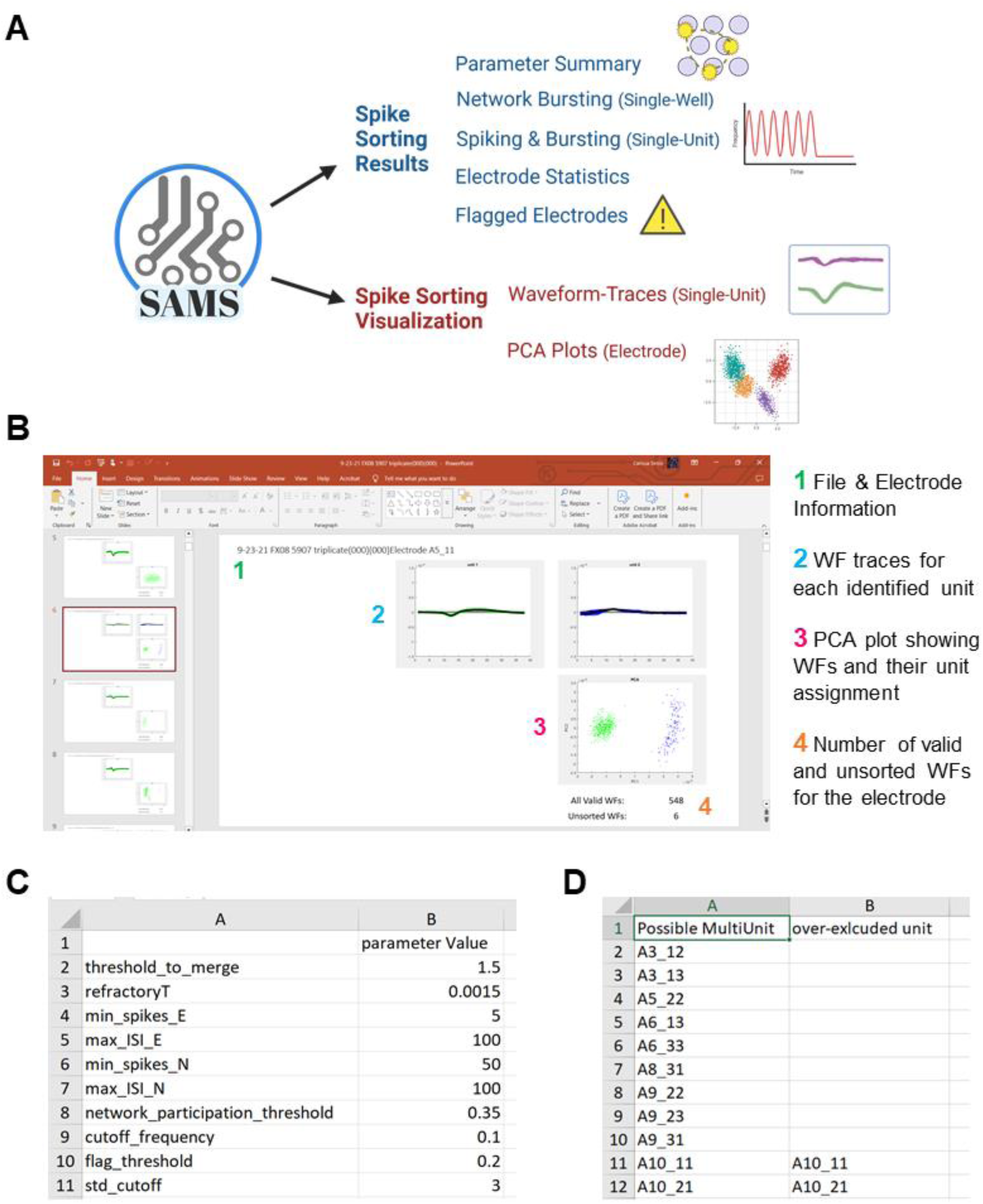
SAMS Output Files for Comprehensive Spike Sorting Analysis. **A.** Explanation of output files and data generated by SAMS. **B.** Electrode-specific spike sorting results are saved in a Powerpoint slide deck. Each slide presents sorting outcomes for an individual electrode (1), displaying color-coded waveforms of defined units (2) alongside corresponding PCA plots (3) and quantification of valid and invalid waveforms (4). This visualization facilitates rapid assessment of automatic sorting quality and aids in identifying potential mis-sorting events. **C.** Parameter list: Summary of the parameter settings used for automated spike sorting analysis. **D.** Quality control: list of potentially mis-sorted electrodes as flagged by SAMS, enabling targeted manual refinement of sorting results.

Updated sorting results matching user-provided curation can then be exported after modifications, closing the verification loop.

### SAMS outperforms commercially available software for spike sorting hPSC-derived neurons

Current recommendations for the spike sorting of MEA data acquired using the Axion Biosystems platform are to use Plexon’s OFS software (**Figure 1A**). We therefore benchmarked the performance of SAMS against various automated spike sorting algorithms offered by OFS. To do this, we first created a “ground truth” spike sorting dataset for comparison, which was achieved by manually sorting previously published MEA data sets recorded from hPSC-derived neurons across multiple parameters and conditions.^31^ OFS was used to manually sort the waveforms and determine the number of units on each electrode. This dataset was used to compare the automated spike sorting performance of both OFS and SAMS (**Figure 4, S2**).

**Figure 4.**
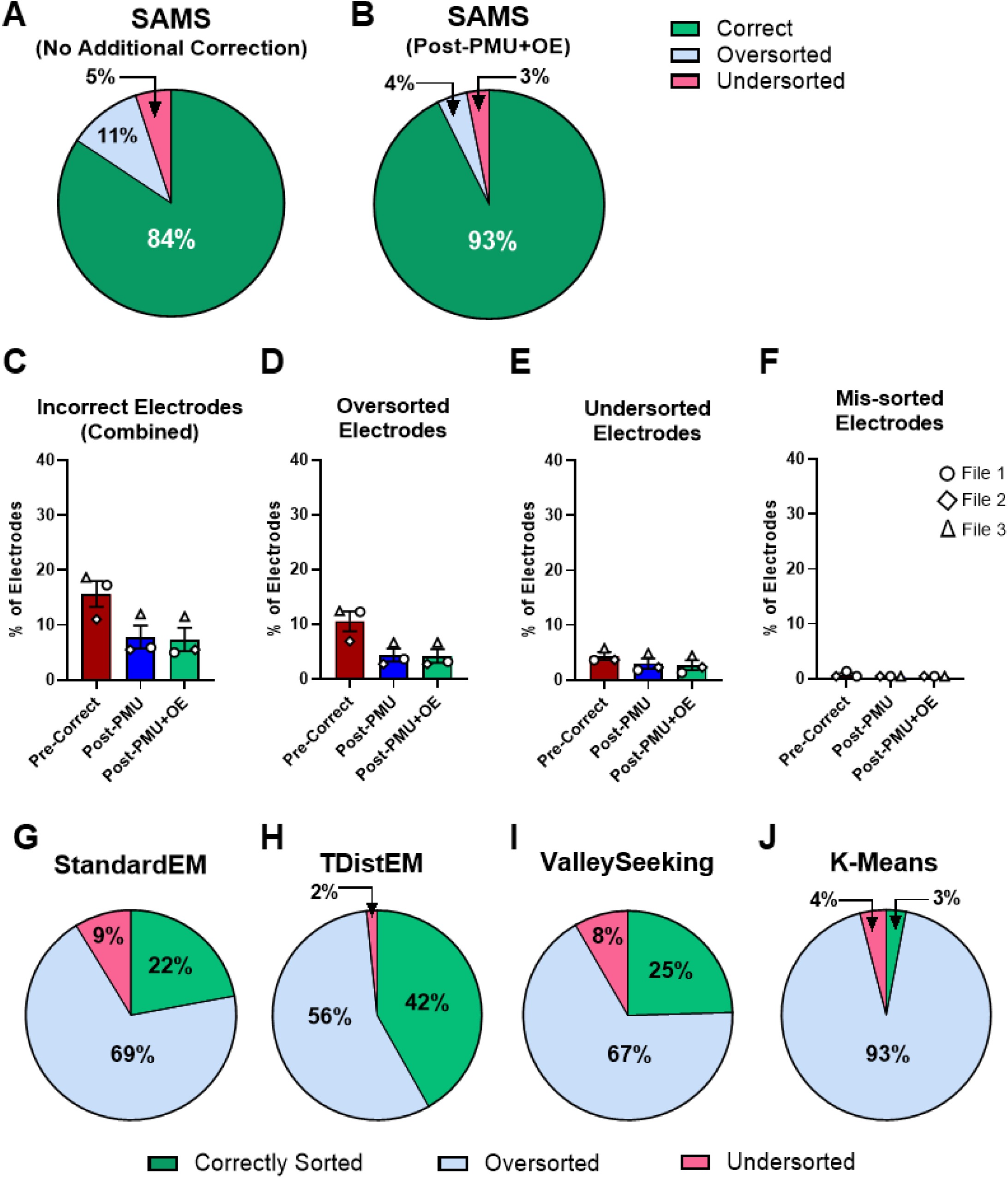
Accuracy of SAMS compared to OFS automated spike sorting algorithms. A-B. Accuracy of SAMS prior to manual correction of any electrodes **(A)**, or after manually correcting any incorrect electrodes flagged by SAMS **(B)**. **C-F.** Percentage of electrodes that were incorrectly sorted by SAMS prior to any manual correction by the user (Pre-Correct, red), after only correcting any incorrect electrodes flagged by SAMS on the “Possible Multi-Unit” list (Post-PMU, blue), or after correction any electrodes on either the “Possible Multi-Unit” list or “Over-Excluded Unit” list of flagged electrodes (Post-PMU+OE, green). **C.** Percentage of total electrodes that were incorrectly sorted. **D.** Percentage of total electrodes that were over-sorted. **E.** Percentage of total electrodes that were under-sorted. **F.** Percentage of total electrodes that were mis-sorted (when the correct number of units was identified but not all waveforms were assigned correctly upon manual inspection). **G-J.** Highest accuracy achieved using automated spike sorting algorithms available in OFS software. **G.** StandardEM: Standard E-M Scan, **H.** TDistEM: T-Dist E-M Scan, **I.** ValleySeeking: Valley Seeking Scan, **J.** KMeans: K-Means Scan. See **Table S3** for settings used and accuracy results for additional analyses performed in OFS. See also **Figure S2**.

We first evaluated the accuracy of the SAMS pipeline alone (without any manual correction of electrodes by the user), as well as the accuracy of SAMS if the user were to manually examine the electrodes flagged by SAMS in the “Manual Check” output lists (**Figure 3D**) and correct any electrodes that were incorrectly sorted. Prior to correction of any flagged electrodes, SAMS correctly sorted more than 84% of electrodes (**Figure 4A).** Furthermore, manual curation of the subset of electrodes identified by SAMS on the “Possible Multi-Unit” flagged electrode list alone (“Post-PMU”), or on both lists of flagged electrodes (“Post-PMU+OE”), further improved this accuracy to 93% on average (**Figure 4B, S2A,B**). Among the less than 20% of incorrectly sorted electrodes (**Figure 4C**), most were errors of over-sorting (6.91-12.44%; **Figure 4D**), while under-sorting (3.64-5.78%, **Figure 4E**) and mis-sorting, when the correct number of units was identified but some of the waveforms were determined to be incorrectly assigned upon manual inspection (0.44-1.36%, **Figure 4F**), were less frequent occurrences.

Next, we compared the accuracy of SAMS to the automated clustering and spike sorting algorithms offered by OFS (**Figure 4G-J, Figure S2**), which were run using default parameters or adjusted parameters (**Table S3**). Of the algorithms available in OFS, Standard EM Scan (**Figure 4G**), T-Dist EM Scan (**Figure 4H**), and Valley Seeking Scan (**Figure 4I**) achieved similar levels of accuracy depending on the file or settings (∼10-42%, **Figure S2C, Table S3**). K-Means Scan, a commonly used algorithm for spike sorting, had even lower accuracy, with fewer than 3% of electrodes correctly sorted (**Figure 4J**). These low accuracy rates were predominantly due over-sorting of waveforms into too many units (**Figure 4G-J, S2D**).

Comparison of the types of errors committed by the OFS algorithms and SAMS indicated similar amount of under-sorting across both platforms (**Figure S2C**), but dramatic improvements by SAMS in the amount of over-sorting errors as compared to OFS (**Figure S2E**). Therefore, SAMS offers significantly improved accuracy in the spike sorting of hPSC-derived neuron MEA data compared to the recommended methods for data acquired using the Axion system, and produces accuracy levels comparable to spike sorting methods used to analyze *in vivo* or simulated MEA datasets.^35–37^

### SAMS is highly accurate in analyzing various recording conditions

MEA recording conditions can vary widely across studies, and MEA data can be influenced by experimental conditions.^38^ Therefore, we next wanted to examine whether the human-machine consistency (HMC) of SAMS was affected by extrinsic experimental factors, or by intrinsic neuronal properties (**Figure 5**).

**Figure 5.**
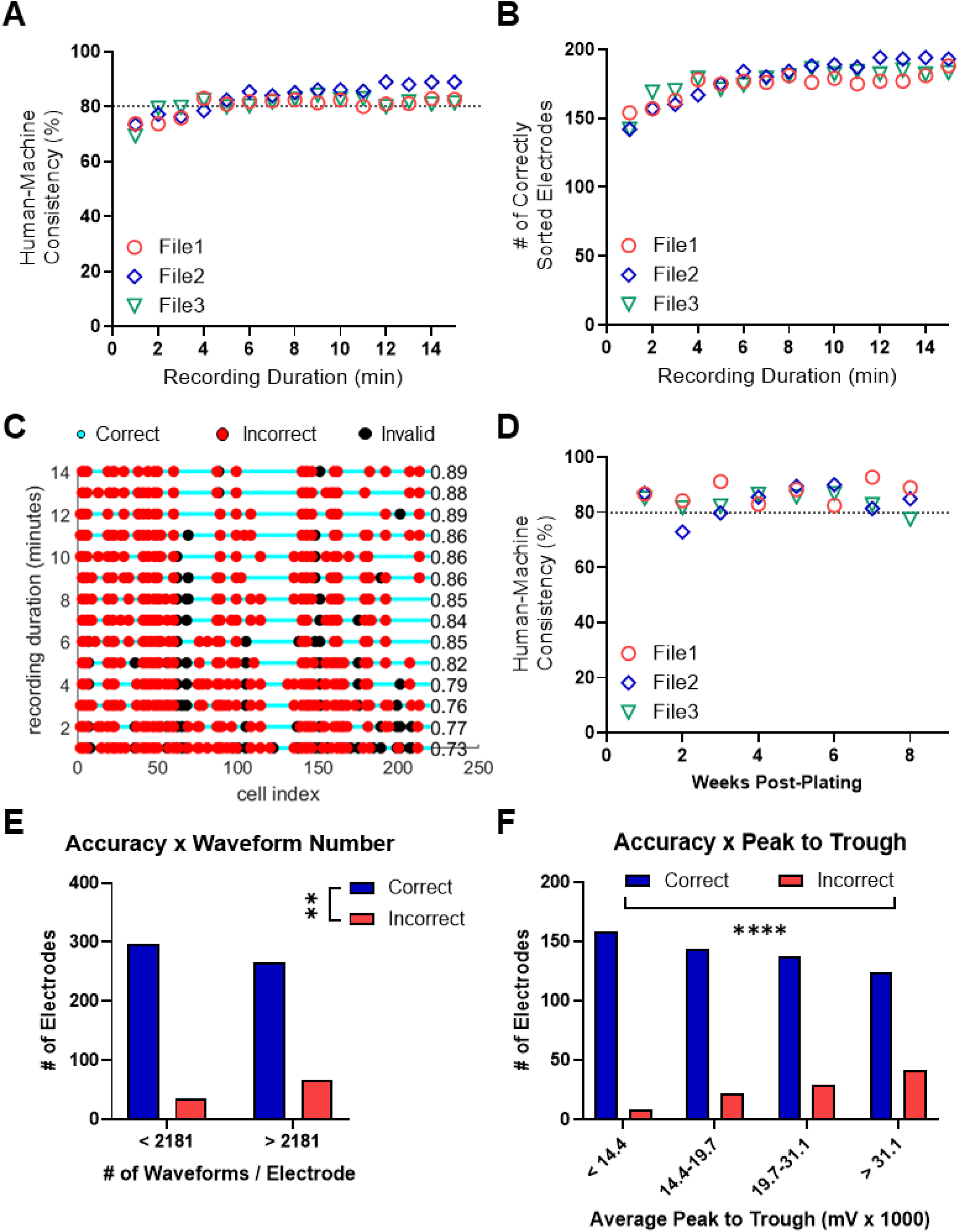
Factors affecting human-machine consistency. **A.** Human-machine consistency over recording duration for three different data files (File 1, 2, 3). Consistency remains above 80% for ≥6 minutes recording durations. **B.** Same data as in A, but shown as the number of correctly sorted electrodes as a function of recording duration, showing an increase in correctly sorted electrodes over time. **C.** Graphical representation of electrode-specific sorting performance (right y-axis) across different recording durations (left y-axis) showing that recording duration does not dramatically impact sorting accuracy on the single-electrode level. Cyan indicates correctly sorted electrodes, red indicates incorrectly sorted electrodes, and black indicates inactive or invalid electrodes. **D.** Human-machine consistency across time of neuronal culturing (1 week to 8 weeks) for three different sets of data files (File 1, 2, 3). Consistency remains stable throughout maturation. **E.** Sorting accuracy relative to the total number of spikes per electrode, split by median spikes per electrode. **F.** Sorting accuracy as a function of average spike amplitude per electrode (mV), split by quartiles of average amplitude. A significant difference in frequency is observed between correctly and incorrectly sorted spikes in both **E** and **F**. See also **Figure S3**. Statistics in E,F: Fisher’s exact test ** p < 0.01; **** p <0.001

Longer recordings generate more data points and increase the analytical burden, while shorter recordings were previously shown to affect MEA data consistency in rat primary neurons.^38^ Therefore, we first assessed whether recording time impacted the performance of SAMS. Specifically, we compared results across different MEA recording durations from 1 to 14 minutes. The HMC (**Figure 5A**) remained high (∼80%) and stable across recording durations of up to 15 minutes for three different files, which contained recordings from six batches of neurons (3 independent batches of differentiation from two hPSC lines). Concurrently, the number of correctly sorted electrodes (**Figure 5B**) increased with recording duration, plateauing around 170-180 electrodes around 10 minutes of recording. **Figure 5C** provides a detailed view of electrode-specific performance across recording durations, revealing patterns of electrodes that were consistently accurately sorted (cyan) compared to our ground truth data set and those requiring manual correction (red). These results reflect the increase in accurately sorted electrodes indicated by the results in **Figure 5B**, and also indicate that certain electrodes are difficult to resolve independent of the recording duration. Because hPSC-derived neurons continue to mature *in vitro* for several weeks or even months after plating,^22,23,30,31,39,40^ we also examined whether the culturing time affected the performance of SAMS. Here, HMC remained stable (75-80%) from 1 week to 8 weeks of *in vitro* culture time (**Figure 5D**). Therefore, SAMS exhibits high accuracy regardless of the developmental stage of the neural network.

We next asked if the accuracy of SAMS was related to the total number of recorded action potentials. Comparison of correctly sorted versus incorrectly sorted electrodes showed significantly fewer total spikes in the correctly sorted compared to the incorrectly sorted electrodes (**Figure S3A**), and there was a significant difference in the frequency of correctly sorted and incorrectly sorted electrodes among electrodes with high and low numbers of waveforms (**Figure 5E**). Therefore, SAMS has good accuracy in sorting low spike count electrodes. We also assessed whether waveform amplitude (peak to trough) affected SAMS accuracy, and found that the amplitude was significantly lower in the correctly-sorted compared to the incorrectly-sorted electrodes (**Figure S3B**), and that there was a significant difference in the frequency of correctly sorted and incorrectly sorted electrodes depending on the waveform amplitude (**Figure 4F**).

These results collectively indicate that SAMS provides robust and accurate spike sorting across various recording conditions and spike characteristics, with particular strengths in handling lower amplitude signals and moderate spike counts. The algorithm’s performance remains consistent across different maturation stages of neuronal cultures, suggesting it will perform well across a wide range of experimental paradigms in electrophysiological studies.

### SAMS unveils electrical activity changes of human neurons with autism-related mutations

To evaluate SAMS as a tool for assessing phenotypic differences among cultured human iPSC-derived neurons, we next analyzed MEA data from two previously published studies from our lab.^30,31^

Since SAMS was developed using neurons derived using a standard dual SMAD-inhibitor-based differentiation protocol,^31,41^ we assessed whether SAMS could also be used to analyze neurons generated through Neurogenin 2 (NGN2) directed differentiation^30^ as these neurons exhibit different temporal trajectories in terms of their differentiation and maturation compared to standard differentiation protocols.^42–44^ We first performed manual spike sorting and calculated the HMC as done above (**Figure 4**). Excitingly, SAMS achieved around 89% accuracy on files analyzed (**Figure 6A**).

**Figure 6.**
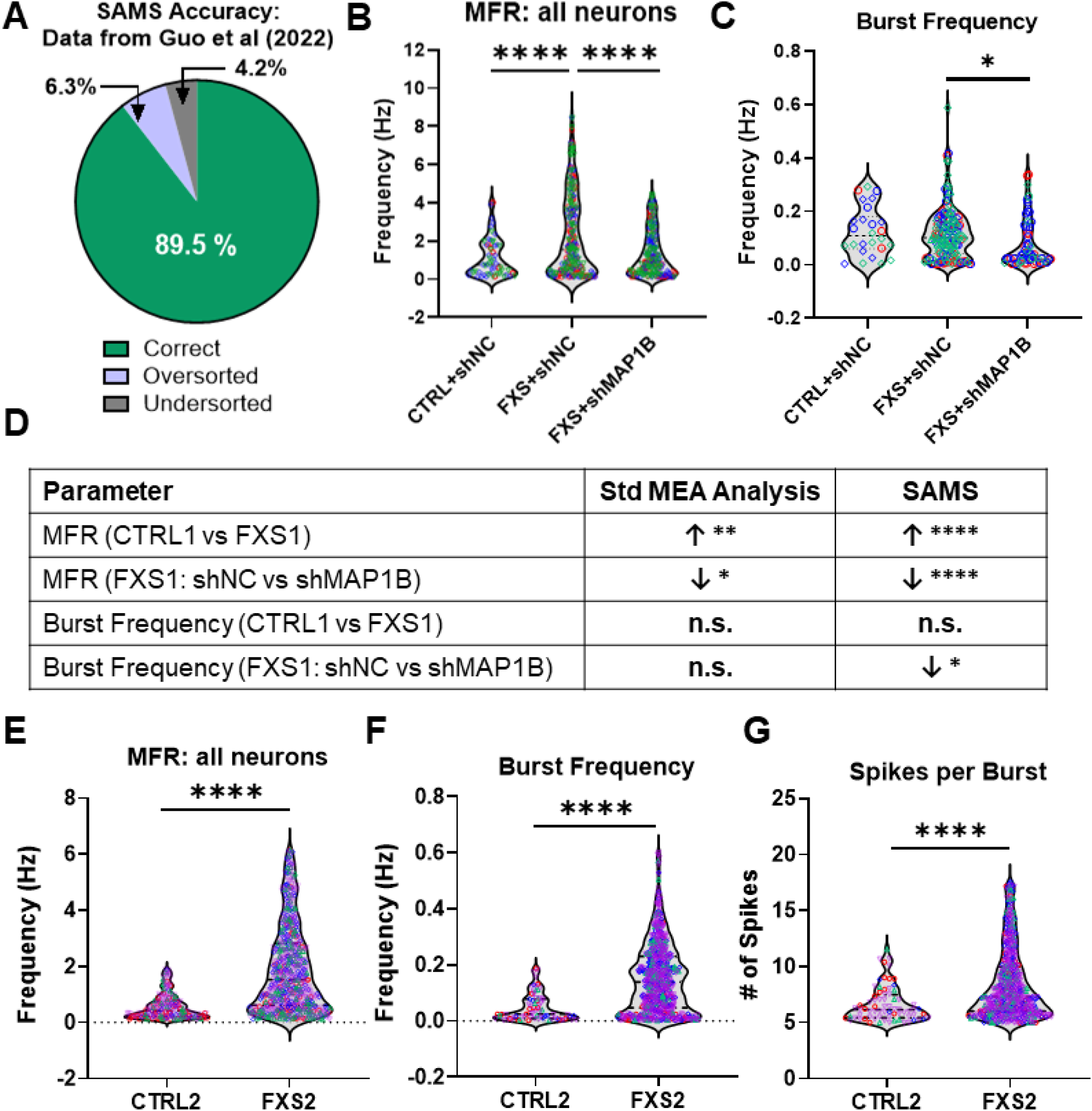
Comparison of SAMS findings to published bulk MEA analysis results. **A.** Accuracy of SAMS in sorting datasets from Guo et al (2023). n = 2 datasets manually checked for accuracy. **B.** Mean firing rate (MFR) and **C.** burst frequency in CTRL1 + shNC, FXS1 + shNC, and FXS1 + shMAP1B neurons after performing spike sorting with SAMS. Data shown are from N = 3 independent batches of neuronal differentiation for each condition (B: n = 56-235 individual neurons; C: n = 26-128 individual neurons). Brown-Forsythe & Welch ANOVA tests followed by Dunnett’s T3 posthoc test. **D.** Summary of results for two parameters reported in Guo et al (2023) and analysis of the same data by SAMS. **E-G.** Results of spike sorting of data from Shen, Sirois, Guo, Li et al (2023). **E.** Mean firing rate, **F.** burst frequency, and **G.** number of spikes per burst in CTRL2 and FXS2 neurons. N = 4 independent batches of neuronal differentiation (E: n = 249-759 neurons; F-G: n = 75 – 627 individual neurons). Welch’s t-test. * p < 0.05, ** p < 0.01, *** p < 0.005, **** p < 0.001 See also **Figure S4**

Using standard Axion MEA analysis without spike sorting, we previously showed that both FXS hPSC-derived neurons and *MAP1B* overexpression (MAP1B-EE) neurons exhibit higher MFR that can be reduced by knocking down *MAP1B* (**Figure 6D, S4A**).^30^ The results following analysis with SAMS yielded consistent findings with our published data, in terms of the directionality of the data trends, as well as whether the differences between groups were statistically significant (**Figure 6B-D; Figure S4A-C**). Importantly, SAMS resolved additional differences among experimental conditions that were missed without spike sorting. Specifically, knocking down *MAP1B* in FXS neurons significantly decreased burst frequency at the single-unit level (**Figure 6C**). Therefore, SAMS not only yielded consistent results as the standard MEA analysis, but also identified additional electrophysiological features that were previously missed.

We next returned to our original data set from Figures 4 and 5, for which SAMS had accurately sorted approximately 84% of the electrodes analyzed (**Figure 4A**), and assessed whether SAMS, without manual correction of any flagged electrodes, was able to yield conclusions consistent with the findings using standard MEA analysis.^31^ We analyzed MEA recordings from four independent batches of neurons generated from a control individual (CTRL2) and an individual with Fragile X Syndrome (FXS2), on which our lab previously performed extensive characterization using MEA.^31^ Excitingly, the differences between CTRL2 and FXS2 neurons observed using standard MEA analysis were consistent with the results generated using SAMS (**Figure 6E-G; Table S4**). SAMS was also able to further resolve differences in burst duration (**Figure S4D-F)**, as well as in other bursting parameters including inter-burst interval (**Table S4**), that did not reach statistical significance using standard MEA analysis methods.

We last wanted to confirm whether sorting of spikes into individual units, and removal of low-activity or outlier waveforms, influenced the electrode level network activity. To do this, the single-unit data for each electrode was combined, and then electrode-level network activity parameters were calculated from this recombined data. We again observed consistent differences in the network activity between FXS2 and CTRL2 neurons as we did with standard MEA analysis (**Figure S4G-I; Table S4**), with the change in statistical significance observed for the network burst duration (**Figure S4I**) being due to performing the analysis at the single-well level. Together, these results indicate that spike sorting analysis using SAMS does not drastically alter the conclusions drawn when making comparisons between groups using standard MEA data analyses.

### SAMS allows for separate analysis of bursting and non-bursting neurons from each electrode

Finally, we wanted to utilize SAMS’ spike sorting capabilities to better understand our neuronal populations. Standard MEA data analysis gives the user an output value that represents the average for a given parameter recorded for each electrode, or across all electrodes in an entire well. For spiking parameters such as mean firing rate, this value reflects all neurons that exhibit action potential firing on a given electrode or in a single well, combining firing rates from neurons that exhibit bursting activity with those neurons that do not. As differences in bursting activity could reflect differences in electrical maturation or other neuronal properties,^41–43^ and bursting differences have been observed in stem cell-derived neurons modeling multiple neurodevelopmental and neuropsychiatric disorders,^30,31,45–48^ spike sorting can enable the user to make direct comparisons of neurons that are, for example, more similar in their electrical maturity.

One robust phenotype of neurons derived from FXS patients that has been observed across multiple laboratories and iPSC lines is that FXS neurons exhibit overall hyperexcitability compared to control neurons, with increased rates of spiking and bursting.^30,31,49–52^ Different genetic or pharmacological interventions to ameliorate these *in vitro* FXS phenotypes decrease overall firing rates but can have varying effects on bursting activity or network activity.^30,31^

The spike sorting results generated by SAMS allowed us to compare bursting and non-bursting neurons in our MEA datasets obtained from iPSC-derived neurons from two FXS and two control individuals (FXS1, FXS2, CTRL1, CTRL2). Neurons derived from both FXS patients exhibited increased firing rates as determined using standard bulk MEA analysis,^30,31^ or when the single unit activity from both bursting and non-bursting neurons was combined into one group (**Figure 6B,E**). FXS neurons that exhibited bursting activity from both iPSC lines had increased firing rates compared to their respective controls (**Figure 7A,D**). Interestingly, the same was not true when we compared firing rates in neurons that did not exhibit bursting activity: non-bursting FXS1 neurons had similar firing rates as the non-bursting CTRL1 neurons, while FXS2 neurons had decreased firing rates compared to CTRL2 neurons (**Figure 7B,E**).

**Figure 7.**
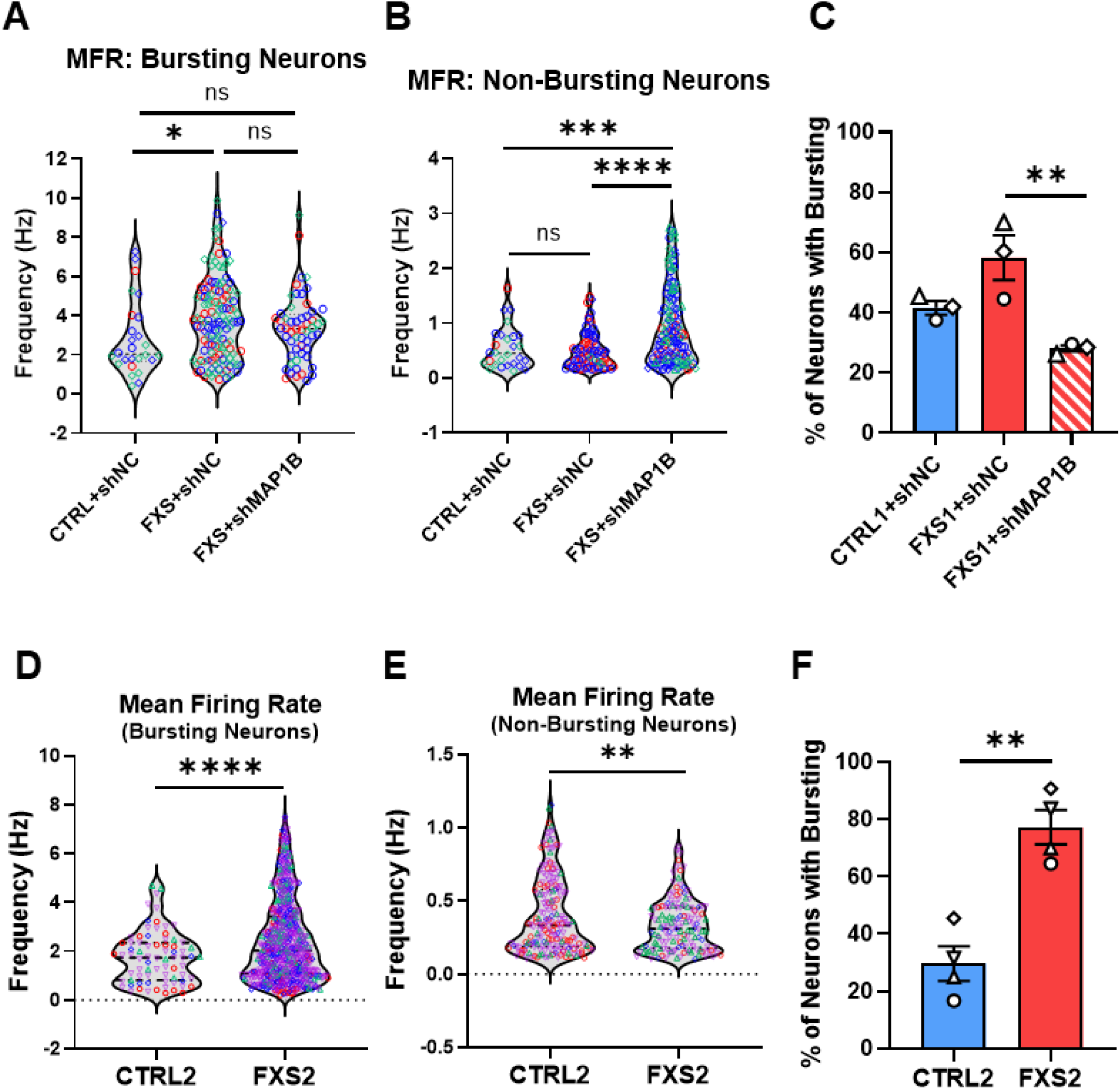
Examination of bursting and non-bursting neuronal populations using SAMS. The same data as in Figure 6, but with separation of neurons based on the presence or absence of bursting activity. **A-C.:** Data from Guo et al (2023). **D-F.** Data from Shen, Sirois, Guo, Li et al (2023). **A,D.** Mean firing rate in neurons that exhibited bursting activity. **B,E.** Mean firing rate in neurons that did not exhibit any bursting activity. **C,F.** Percentage of neurons in each batch that exhibited any bursting activity. Data shown in (A-C) are from N = 3 independent batches of neuronal differentiation, while data from (D-F) are from N = 4 independent batches of neuronal differentiation. Statistics: A,B: Brown-Forsythe & Welch ANOVA tests followed by Dunnett’s T3 posthoc test (n = 26 – 165 individual neurons); C: ordinary one-way ANOVA followed by Tukey’s posthoc test; D,E: Welch’s t-test (n = 90 – 652 individual neurons); F: unpaired Student’s t-test.

Additionally, while the proportion of neurons that exhibited bursting activity was higher in both FXS lines, this difference was significant only in FXS2 neurons (**Figure 7C,F**).

Finally, we wondered whether these neuronal subpopulations after spike sorting might reveal additional mechanistic insights into our previously published finding that shRNA knockdown of the *MAP1B* gene in FXS1 neurons (“FXS1/shMAP1B”) could partially rescue FXS1 MEA phenotypes when analyzed using standard methods.^30^ Spike sorting revealed that knockdown of *MAP1B* in FXS1 neurons did not change the mean firing rate of bursting FXS1 neurons (**Figure 7A**), but actually increased the mean firing rate of non-bursting FXS1 neurons (**Figure 7B**) and decreased the proportion of FXS1 neurons that exhibited any bursting activity (**Figure 7C**). Together, these results demonstrate the utility of using SAMS to perform spike sorting and also illustrate that additional results can be obtained by performing spike sorting of MEA data from human iPSC-derived neurons, which would otherwise be missed by analyzing data using the standard methods.

## DISCUSSION

SAMS represents a significant advancement in the field of spike sorting for low-density MEA recordings, particularly for those involving hPSC-derived neurons. By integrating multiple algorithms and quality control measures, SAMS addresses several key challenges in the current spike sorting landscape. First, SAMS significantly streamlines the spike sorting workflow for MEA data acquired using the Axion MEA platform. The traditional pipeline, involving multiple software platforms and file formats, and extensive manual interventions, is time-consuming and requires specialized coding skills. SAMS consolidates these steps into a single, user-friendly interface, making advanced spike sorting techniques accessible to a broader range of researchers. This democratization of spike sorting technology has the potential to accelerate neuroscience research, particularly in the field of *in vitro* disease modeling using hPSC-derived neurons.

The performance of SAMS is superior to the automated sorting algorithms in the OFS software that is recommended to use with Axion MEA data. Our analysis shows that SAMS achieves high human-machine consistency (>80%) across various recording durations, neuronal culture maturities, batches of neurons, and neurons from different disease conditions. These levels of accuracy are similar to algorithms that have been developed for spike sorting of MEA data acquired from other platforms or tissue types.^24,35–37^ This robustness is crucial for ensuring reliable results across diverse experimental paradigms. The algorithm’s ability to maintain performance across different maturation stages of neuronal cultures (1-8 weeks) is particularly valuable for developmental studies, long-term experiments, and repeated recordings of the same neurons over weeks in culture. SAMS also demonstrates specific strengths in handling challenging spike characteristics. It performs significantly better with lower amplitude spikes and smaller peak-to-trough amplitudes, areas where traditional sorting methods can struggle.

The multi-step approach employed by SAMS – combining spectral clustering, Hartigan’s Dip Test, Dynamic Time Warping, and waveform validation – provides a comprehensive solution to common spike sorting issues by compensating for the limitations of individual spike sorting algorithms. This approach effectively addresses both under-sorting and over-sorting problems, maintaining a balance between sensitivity and specificity in unit identification. The levels of accuracy achieved by SAMS, particularly after validation of the Flagged Electrodes lists, is particularly significant in the context of high-content screening using MEA plates in 48-or 96-well plate formats, as these features dramatically reduce the amount of time the user will need to spend checking electrodes before they obtain their final results. The application of SAMS to existing MEA datasets demonstrates its practical utility in research scenarios such as disease modeling or gene knockdown studies. This versatility makes SAMS a valuable tool for a wide range of neuroscience applications, from basic research to disease modeling and drug discovery.

In conclusion, SAMS represents a significant step forward in making sophisticated spike sorting accessible and efficient for researchers working with low-density MEAs and stem cell-derived neurons. By reducing the technical barriers to high-quality spike sorting, SAMS has the potential to accelerate research in critical areas such as neurodevelopmental disorders, including FXS. As the field of neuroscience continues to generate increasingly complex datasets, tools like SAMS will play a crucial role in extracting meaningful insights from neural recordings, ultimately contributing to our understanding of brain function and dysfunction.

### Limitations of this study

It is important to note that SAMS, like other systems, has limitations. The algorithm’s performance appears to decrease slightly with very high spike counts per electrode, suggesting potential room for improvement in handling highly active neurons or high-noise scenarios.

Future iterations of SAMS may incorporate machine learning techniques to further refine its performance in these edge cases. Additionally, this current iteration of SAMS cannot identify or define network bursts at the single unit level, therefore network activity data currently still needs to be examined on an electrode-to-electrode basis. This version of SAMS also only reports electrode-level network spiking and bursting metrics, therefore measures of synchrony would still need to be acquired through standard bulk MEA analysis. Finally, while we were able to validate SAMS using the data files acquired by independent investigators in our group over multiple years, we were unable to access recordings of similar durations from other investigators to test SAMS pipeline due to a lack of data availability. The data files used for the development of SAMS also almost exclusively consisted of cortical excitatory neurons, so analysis of recordings containing purely inhibitory or other neuronal subtypes may produce differences in the overall accuracy. Therefore, as SAMS is applied to a broader variety data sets, additional updates and refinements to the SAMS algorithm can be made to retain its high levels of accuracy.

## ACKNOWLEDGEMENTS

We thank the Zhao lab for helpful discussion. This work was supported by the DOD IIRA grant W81XWH-22-1-0621 (to X.Z.), National Institutes of Health (R01MH118827, R01MH116582, R01MH136152, and R01NS138268 to X.Z.; R01EY029438 and R01EY035005 to A.R.; P50HD105353 to Waisman Center), Kellett Mid-Career Award, Vilas Distinguished Achievement Professorship, Wisconsin Alumni Research Foundation, Jenni and Kyle Professorship, and Eagle Autism Foundation (to X.Z.), Simons Foundation Autism Research Initiative pilot grant (to X.Z.), postdoctoral fellowships from the University of Wisconsin Stem Cell and Regenerative Medicine Center, FRAXA, and the Autism Science Foundation (to C.L.S.). This work was also supported by SciMed scholarships, T32 GM141013 Molecular Pharmacology training grant and predoctoral fellowship from the Wisconsin Stem Cell and Regeneration Medicine Center (to N.M.M-A), and the National Institute of Neurological Disorders and Stroke and the Office of the Director’s Common Fund at the National Institutes of Health (grant DP2NS122605 to A.H.).

## AUTHOR CONTRIBUTIONS

C.L.S. and X.Z. conceived the concept and designed the study. X.R. developed the method with help from A.R., R.D., and A.H. C.S. and X.R. performed accuracy checking, created figures and tables. N. M-A created the logo. X.R., C.L.S. and X.Z. wrote the manuscript. All authors edited the manuscript.

## DECLARATION OF INTERESTS

The authors declare no competing interests.

## RESOURCE AVAILABILITY

The code and files used for generating the figures in this paper can be accessed at https://github.com/xiaoxuanren/SAMS-. All other data are available in the main article or in the supplemental information. Further information and requests for resources should be directed to the corresponding authors.

## SUPPLEMENTAL INFORMATION

Supplemental Figures S1-S4. Supplemental Tables S1-S5.

## METHODS

### SAMS

The SAMS pipeline employs a multi-step approach to achieve robust spike sorting for *in vitro* MEA recordings of hPSC-derived neurons, the key components of which are outlined below:

#### Clustering Methods

Initial unit assignments are performed using spectral clustering,^32^ an unsupervised technique that analyzes the eigenvalues of the similarity matrix of the waveform data. The algorithm solves the following eigenvalue problem:

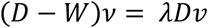

Where *D* is the degree matrix, *W* is the weighted adjacency matrix, *v* is the eigenvector, and λ is the eigenvalue. The eigenvectors corresponding to the k smallest eigenvalues are used to partition the data into k clusters. This method maximizes between-cluster variance while minimizing within-cluster variance, providing a solid foundation for subsequent refinement steps.

#### Hartigan’s Dip Test (HDT)

The Hartigan’s Dip Test^33^ was implemented to assess the initial clustering quality. The dip statistic D is defined as:

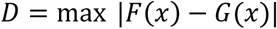

Where *F*(*x*) is the empirical distribution function and *G*(*x*) is the unimodal distribution function that minimizes the maximum difference. Clusters that fail the dip test (i.e., *D* exceeds a critical value, here, 0.1), indicating multimodality, are recursively split to improve sorting accuracy. This step helps identify and separate potentially overlapping units that may have been merged in the initial clustering (**Figure 1C,D**).

#### Dynamic Time Warping (DTW) Distance

Following the cluster refinement, we address potential over-sorting errors using Dynamic Time Warping (see **Figures 1E, 1F, and S1**).^34^ The DTW distance between two time series X = (x_1_,…, x_n_) and Y = (y_1_,…, y_m_) is calculated as:

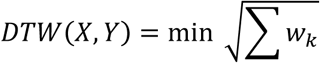

Where *w_k_* is the k-th element of a warping path W. The optimal warping path is found using dynamic programming to minimize the cumulative distance between the aligned elements of two waveforms: X and Y. Units with DTW distances below a specified threshold (here: 1.5) are merged, helping to consolidate spikes from the same neuron that may have been incorrectly split due to temporal variations in their waveforms.

#### Excluding Invalid Waveforms

In the final stage of our pipeline, we implement rigorous quality control measures to ensure the validity of sorted units. Waveforms are excluded if they deviate significantly from each unit’s mean template μ:

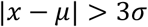

Where *x* is a waveform, μ is the mean template of the unit, and σ is the standard deviation of the unit’s waveforms. Additionally, we discard units with firing rates (fr) below a threshold θ: *f*_r_ < θ, specified by the user (here θ = 0.1 Hz) and have an inter-spike interval (ISI) below a threshold (here: 1.5 ms). This conservative approach helps maintain the integrity of the sorted data for further analyses (see **Figure 1G,H**).

#### Criteria for Flagging of Potentially Mis-sorted Units

SAMS implements two categories of flagged electrodes: Possible Multi-Unit (PMU) and Over-Excluded Unit (OEU). These flags help identify potential issues in spike sorting for further review.

(1) Possible Multi-Unit (PMU) Flagging

An electrode may be flagged as PMU under two conditions:

- Hartigan’s Dip Test Failure: If an electrode fails the Hartigan’s dip test, indicating a single unit may actually be two units, we examine the overlap of the waveform ranges. If the overlap exceeds 70%, the original unit is not split, but the electrode is flagged as PMU.
- Dynamic Time Warping (DTW) Distance: We calculate the DTW distance between aligned elements of two waveforms. Units with DTW distances below a specified threshold (default 1.5) are flagged as PMU. This threshold was determined through expert analysis of thousands of electrodes across multiple files, where a DTW distance around 1.2 typically indicated over-classification (see **Fig. S1**).

(2) Over-Excluded Unit (OEU) Flagging

An electrode is flagged as OEU if the total number of excluded waveforms during the invalid waveform exclusion procedure exceeds a certain threshold (default: 10% of total waveforms).

Both the DTW distance threshold for PMU and the exclusion threshold for OEU can be adjusted by users in the SAMS interface to suit specific experimental needs.

### Generation of aligned Axion spike files for spike sorting

To generate annotated spike files from previously recorded raw MEA data files, raw files were loaded into the AxIS navigator software (Axion Biosystems), then the “Spontaneous” configuration was selected. The Spike Detector was set to “Peak Adaptive Threshold”, but all other settings for the Spike Detector, as well as all settings for the Burst Detector and Statistics Compilier, were set at the default values. The “Peak Adaptive Threshold” setting allows AxIS Navigator to generate spike files that contain annotated spikes and burst information, but also aligns the waveforms to facilitate spike sorting. The specific values for each of the Spike Detector settings is detailed in **Table S6**. Each spike file was then converted to the Neuroexplorer format using Axion’s Data Export Tool software so it was compatible with Plexon’s Offline Sorter (OFS) software.

### Manual (user) spike sorting of MEA data files

To assess the accuracy (human machine consistency (HMC)) of both SAMS and OFS spike sorting algorithms, the data files were first analyzed by hand to determine the number of units on each electrode in each data file. Each converted spike file was loaded into OFS to enable viewing of unsorted aligned waveform shapes and waveform PCA data, and manual sorting of waveforms into putative units. Manual sorting of waveforms was performed as follows: First, PCA clustering and the peaks of aligned waveforms were used to define putative units. Then each putative unit was subdivided into smaller subunits containing 50 to 200 waveforms on average. These subunits were manually inspected to compare their similarity, and two subunits were considered to be from the same unit based on the waveform amplitude, waveform shape similarity, alignment of peaks and alignment of troughs. Units with less than 0.1 Hz were excluded.^21,23^

### SAMS and OFS Accuracy Comparisons

To assess the accuracy of SAMS and OFS algorithms, the number of units identified on each electrode was compared to the number of units identified by manual spike sorting. SAMS was run using the default settings (see **Table S2**). Four different automated spike sorting algorithms available in OFS software were tested using default and adjusted parameters, the details of which are provided in **Table S3**. An electrode was classified as incorrectly “over-sorted” when the number of units identified by an automated spike sorting algorithm was greater than the number of units identified by manual spike sorting. Similarly, an electrode was considered incorrectly “under-sorted” when the number of units identified by an automated spike sorting algorithm was less than the number of units identified by manual spike sorting.

To further assess the accuracy of SAMS, the PCA plots and waveform plots for each electrode were manually checked to confirm whether the waveforms appeared to be correctly assigned to the identified units. Although this was uncommon (see **Figure 4F**), an electrode was classified as incorrectly “mis-sorted” when the number of units identified by SAMS matched the number of units identified by manual spike sorting, but when the assignment of waveforms was incorrect upon user inspection.

We analyzed the performance of SAMS on three different files (File 1, File 2, and File 3) across various recording durations and maturation stages. We first assessed the algorithm’s performance on the full recording length (15 minutes) for each file, establishing a baseline for comparison. We then evaluated the algorithm’s performance at different recording durations, ranging from 1 to 14 minutes, in one-minute increments (see **Figure 4A,B**), and across different maturation stages, from 1 to 8 weeks (see **Figure 4D**).

### MEA Data sets

All MEA datasets from hPSC-derived neurons used in this study were previously published.^30,31^ Data from Shen, Sirois, Guo, Li et al. (2023) were acquired from neurons differentiated using a standard dual SMAD-inhibition monolayer-based protocol, in which neural progenitor cells (NPCs) were plated for terminal differentiation on MEA plates, then co-cultured with mouse primary astrocytes that were added one week after neuron plating.^31^ Neurons were recorded weekly, starting at one week after the addition of astrocytes, for a total of 8 weeks, for 15 minutes in 5% CO_2_ at 37°C. The labels “FXS2” and “CTRL2” to refer to data from the WC053i-FX08-25 and WC031i-5907-6 lines, respectively (for more information, refer to Shen, Sirois, Guo, Li et al, 2023). Data from Guo et al (2022) were acquired from neurons differentiated using an NGN2-based protocol, in which NPCs were co-transduced with lentiviruses containing TetO-NGN2-Neo and reverse tetracycline-controlled transactivator (rtTA) then treated with doxycycline and selected using G418.^30^ The resulting cells were plated onto MEA plates, then co-cultured with mouse primary astrocytes that were added one week after neuron plating. Neurons were recorded weekly, starting at one week after the addition of astrocytes, for a total of 6 weeks, for 10 minutes in 5% CO_2_ at 37°C. In this paper, we used the labels “FXS1” to refer to the WC007i-FX13-2 line, “CTRL1” to refer to the WC032i-6007-1 line, “CTRL3” to refer to idCas9A-H9 infected with *LV-sgCtrl* and “MAP1B-EE” to refer to idCas9A-H9 infected with *LV-sgMAP1B* (for more information, please refer to Guo et al).

## DATA ANALYSIS AND STATISTICS

The electrode visualization diagram in **Figure 5C** was generated using Matlab. For all other figures, graphing and statistical analysis was performed using Prism software (Graphpad). Prior to testing for significant differences, outliers were identified using the ROUT method (Q = 1%) and then removed. For comparisons between two conditions, unpaired, two-tailed student’s t-test (no significant difference in variance between conditions) or two-tailed, unpaired t-test with Welch’s correction (significant difference in variance between conditions) was performed. For comparisons between more than two conditions, ordinary one-way ANOVA followed by Tukey’s posthoc test (no significant difference in variance between conditions), or Brown-Forsythe and Welch’s ANOVA tests followed by Dunnett’s T3 posthoc test (significant difference in variance between conditions) was performed. For these tests, comparisons were performed by treating each individual neuron as a single data point, with the exception of Figures S4E (N = 4 batches of neurons), S4F (n = 41-54 wells), S4J (N = 3-4 batches of neurons), S4K (n = 12-49 wells), 7C (N = 3 batches of neurons), and 7F (N = 4 batches of neurons), which batches or wells were used for comparison.

For Figure 5E-F, Fisher’s exact test was performed to test for significant differences in distribution between groups. The number of wells and electrodes analyzed for each condition and data file are contained in **Supplemental Table S5.**

## Notes

### Competing Interest Statement

The authors have declared no competing interest.

