## Supplemental Figures and Tables for "A Semi-Automated MEA Spike sorting (SAMS) method for high throughput assessment of cultured neurons"

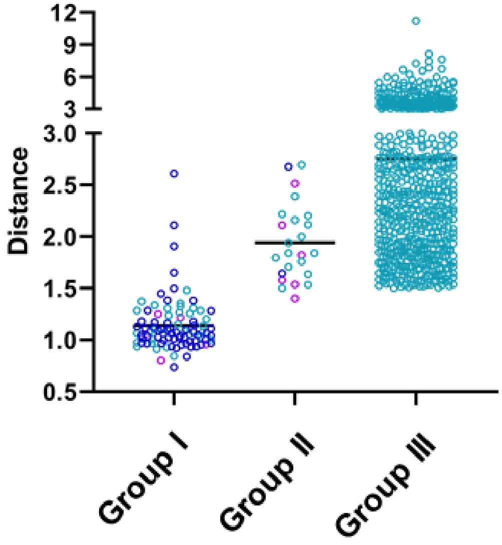


**Figure S1. Selection of default value DTW threshold used in SAMS.**

DTW distances for three groups of sorted spikes: Group I: units that were over-sorted (mean DTW distance ~ 1.2). Group II: units with ambiguous classifications (mean DTW distance ~ 2.0). These cases represent waveforms classified as originating from two different units, but with uncertainty about whether they truly represent distinct units. Group III: Correctly classified units (DTW distance > 1.2). Group III: units for which waveforms were confidently classified as originating from two different cells (DTW distance > 1.5.

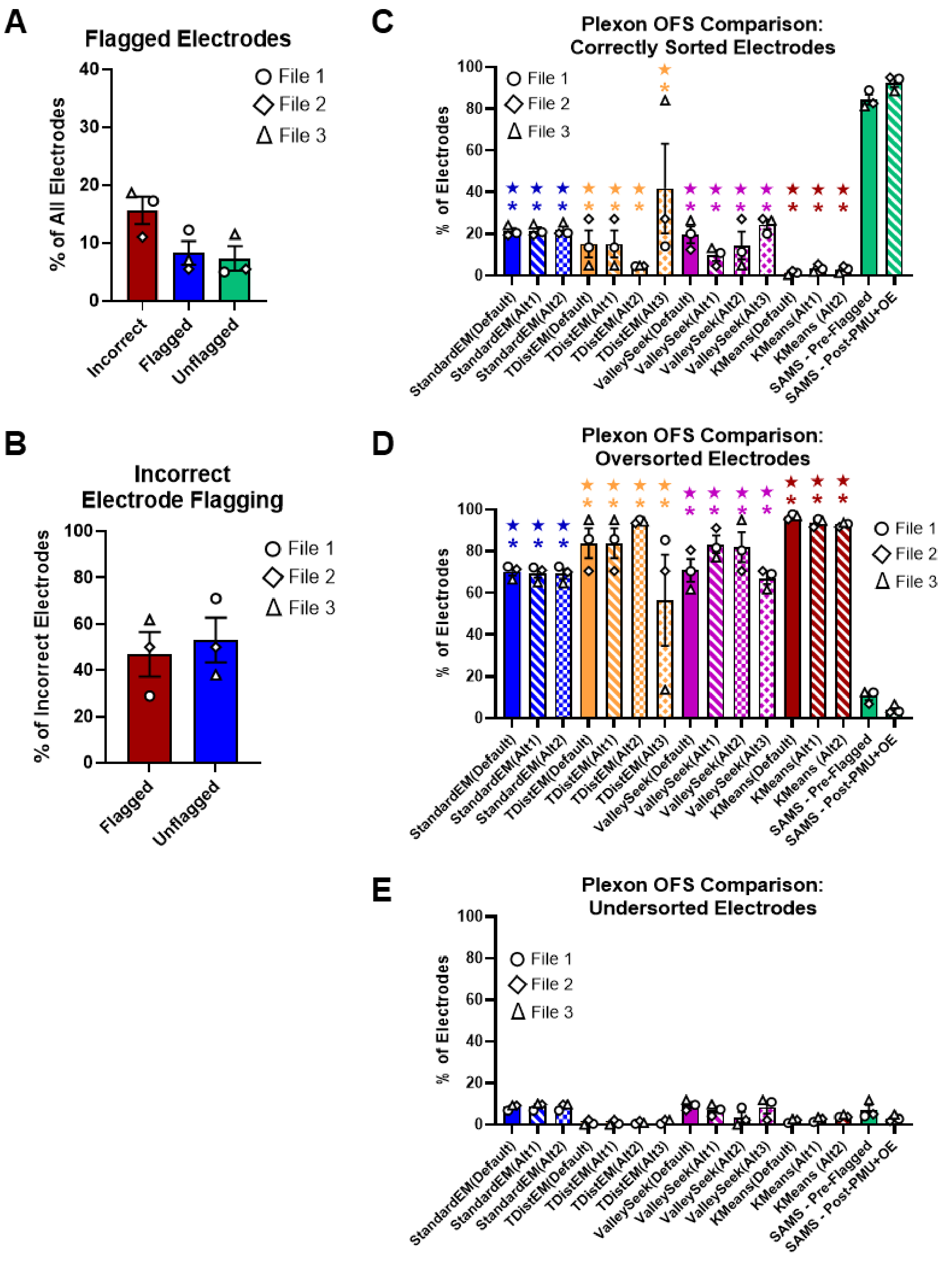

A
Flagged Electrodes
% of All Electrodes
File 1
File 2
File 3
Incorrect
Flagged
Unflagged
B
Incorrect Electrode Flagging
% of Incorrect Electrodes
Flagged
Unflagged
C
Plexon OFS Comparison: Correctly Sorted Electrodes
% of Electrodes
D
Plexon OFS Comparison: Oversorted Electrodes
E
Plexon OFS Comparison: Undersorted Electrodes
StandardEM(Default)
StandardEM(Alt1)
StandardEM(Alt2)
TDistEM(Default)
TDistEM(Alt1)
TDistEM(Alt2)
TDistEM(Alt3)
ValleySeek(Default)
ValleySeek(Alt1)
ValleySeek(Alt2)
ValleySeek(Alt3)
KMeans(Default)
KMeans(Alt1)
KMeans (Alt2)
SAMS - Pre-Flagged
SAMS - Post-PMU+OE

**Figure S2. Comparison of SAMS accuracy with automatic spike sorting algorithms in OFS. (Related to Figure 4)**

**A.** Incorrect electrodes not flagged by SAMS, shown as a proportion of all electrodes. **B.** The same data as in A, but shown as a proportion of incorrect electrodes. **C.** Percentage of all electrodes that were accurately sorted by the indicated algorithm. **D,E.** Percentage of all electrodes that were over-sorted (**D**) or under-sorted (**E**) by the indicated algorithm. Data shown are the same as in **Figure 5** and **Table S3**, which are from N = 3 independent sets of MEA recordings & batches of neurons. Statistics: ordinary one-way ANOVA followed by Tukey's posthoc test; A,B: ★ significant difference when compared to SAMS – Pre-Flagged **** $p < 0.001$; ★ significant difference when compared to SAMS – Post-Flagged **** $p < 0.001$; C: * $p < 0.05$, ** $p < 0.01$; all comparisons with the SAMS-Post Flagged condition were not significant

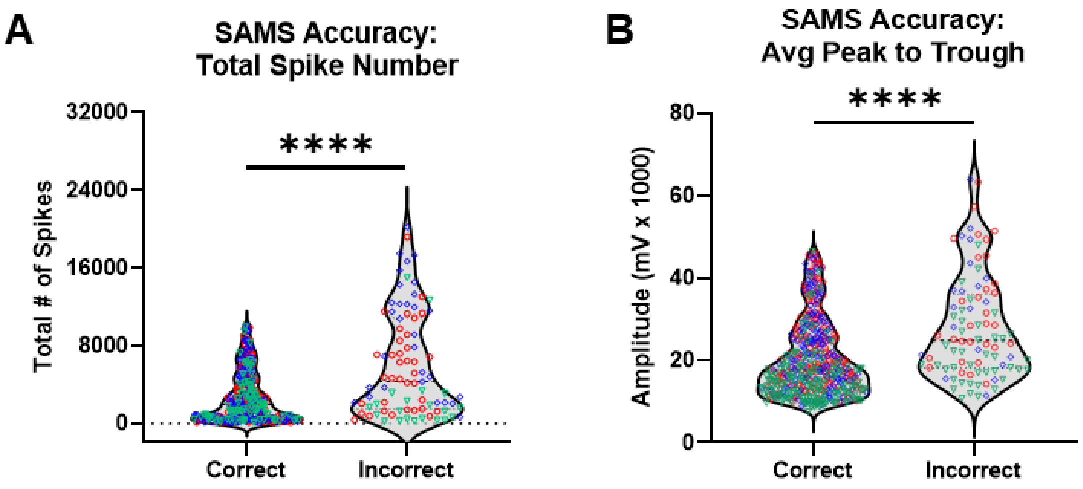


**Figure S3. Factors affecting SAMS accuracy (related to Figure 5)**
**A.** Total number of spikes on each correctly sorted or incorrectly sorted electrode (related to **Figure 4E**). **B.** Average waveform amplitude (peak to trough) of each correctly sorted or incorrectly sorted electrode ( see **Figure 4F**). Statistics: A,B: t-test with Welch's correction, **** $p < 0.001$, n = 105-508 individual neurons from N = 3 independent sets of MEA recordings & batches of neurons.

A

| CTRL3 vs MAP1B-EE | | |
|---|---|---|
| Parameter | Std MEA Analysis | SAMS |
| MFR | ↑ * | n.s. |
| # of Bursts | ↑ ** | ↑ **** |
| Burst Freq | ↑ ** | ↑ **** |
| Burst Duration | n.s. | n.s. |

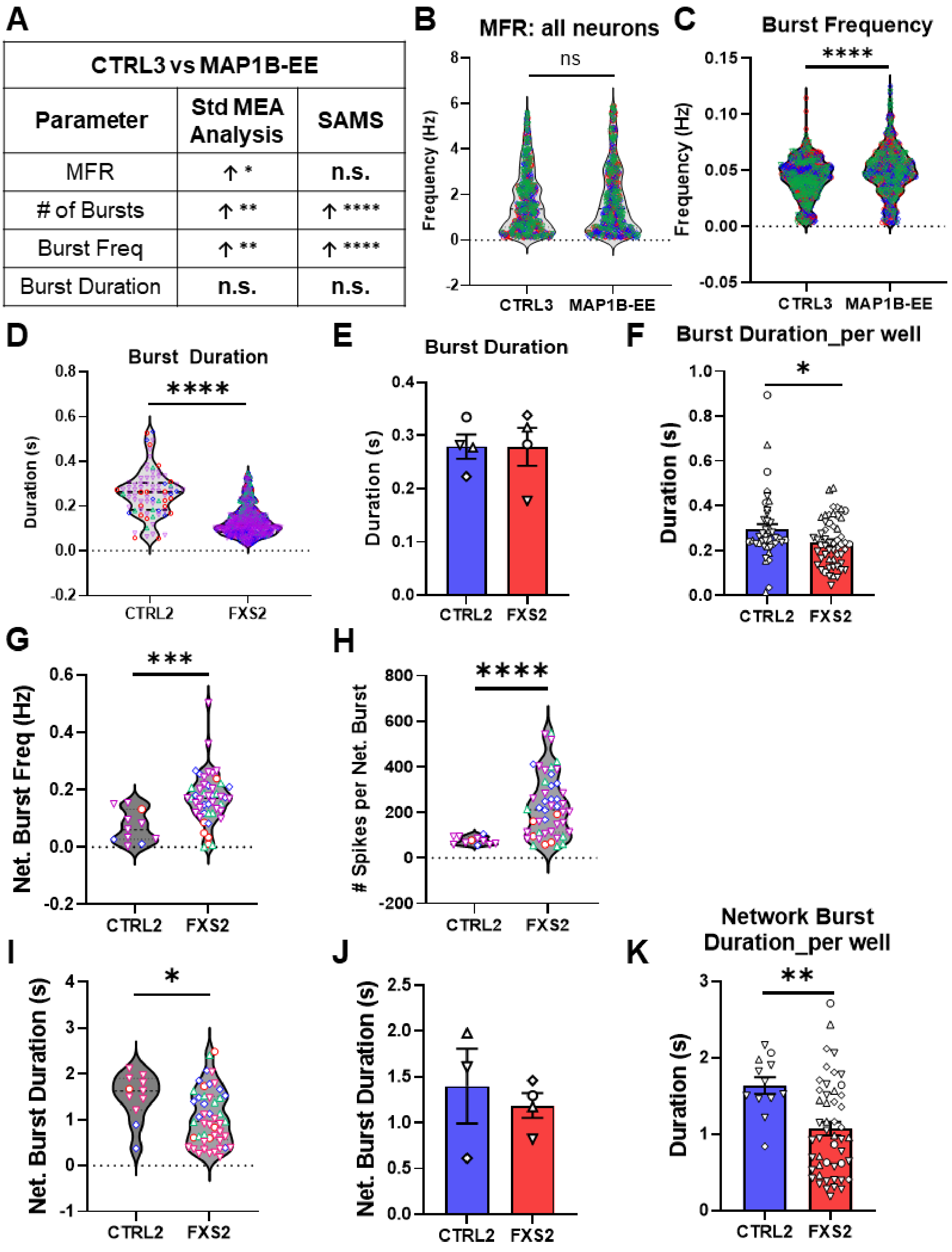

**Figure S4. MEA parameters following analysis using SAMS.**

**A-C.** Analysis of data from Guo et al (2023) comparing control (CTRL3) neurons to neurons with induced expression of *MAP1B* (MAP1B-EE). **A.** Comparison of results using standard MEA analysis (previously published) and following spike sorting using SAMS. **B.** Mean firing rate of all neurons identified using spike sorting. **C.** Burst frequency in neurons identified using spike sorting. Statistics: N = 3 independent batches of differentiation (n = 522 – 549 individual neurons) Welch's t-test *** $p < 0.005$. **D-K.** Analysis of data from Shen, Sirois, Guo, Li, et al (2023) comparing control (CTRL2) and Fragile X Syndrome (FXS2) neurons. **D-F.** Example of a parameter which was inconsistent between SAMS and standard MEA analysis. Comparison of burst duration in individual units after spike sorting with SAMS (**D**) shows decreased burst duration in FXS2 neurons, which was not seen when performing standard MEA analysis (**E**). This difference is likely due to an artifact created by averaging all cells and wells together for each batch of cells, which is supported by comparison of burst duration in individual wells (**F**). **G,H.** Electrode-level network burst parameters following re-merging of spike-sorted data after SAMS: network burst frequency (**G**) and number of spikes per network burst (**H**). **I-J.** Example of a network burst parameter which was inconsistent between SAMS and standard MEA analysis. Comparison of network burst duration in indivsidual units after spike sorting with SAMS (**I**) shows decreased network burst duration in FXS2 neurons, which was not seen when performing standard MEA analysis (**J**) This difference is likely due to an artifact created by averaging all cells and wells together for each batch of cells, which is supported by comparison of network burst duration in individual wells (**K**). Statistics: N = 4 independent batches of neuronal differentiation (D: n = 95-637 individual neurons; F: n = 41-54 individual wells; G-I: n = 12-49 individual neurons; K: n = 12-49 individual wells); D,H: Welch's t-test; E-G,I-K: unpaired student's t-test *$p < 0.05$; ** $p < 0.01$, *** $p < 0.005$; **** $p < 0.001$. See also **Table S4.**

**Table S1. Summary of Available Spike Sorting Programs**

| Name | MEA Type | Recording Environment | References |
|---|---|---|---|
| Klusta suite | High density | *in vivo* | Rossant (2016)[1] |
| Mountainsort4 | Low or high density | *in vivo* | Chung (2017)[2] |
| Ironclust | High density | *in vivo* | Jun (2017) *bioRxiv preprint*[3] *[Benchmarked in Magland (2020)]*[4] |
| WaveClus | Low density | *in vivo* | Quiroga (2004);[5] Chaure (2018)[6] |
| YASS | High density | *in vivo* | Lee (2020) *bioRxiv preprint*[7] |
| HTsort | High density | *in vivo* | Chen (2021)[8] |
| Kilosort4 | High density | *in vivo* | Pachitariu (2024)[9] |
| GEMsort | High density | *in vivo* | Mohammadi (2024)[10] |
| MixFMM | Low or high density | *in vivo* | Rueda & Rodriguez-Collado (2023)[11] |
| MultiFq | Low density | *in vivo* | Wang (2023)[12] |
| Full Binary Pursuit (FBP) | Low or high density | *in vivo* | Hall (2021)[13] |
| SpyKING CIRCUS | High density | *in vivo, ex vivo* | Yger (2018)[14] |
| Offline Sorter (Plexon) | Low density | *in vivo, in vitro* | https://plexon.com/products/offline-sorter/ |
| HerdingSpikes2 | High density | *ex vivo* | Hilgen et al (2017)[15] |
| *[not named]* | High density | *ex vivo* | Diggelmann (2018)[16] |
| *[not named]* | Low density | *in vitro (rat - cortical)* | Negri (2020)[17] |
| MEA-ToolBox | Low or high density | *in vitro* | Hu (2022)[18] |
| μSpikeHunter | High density | *in vitro (rat – cortical; mouse – DRG)* | Heiney (2019)[19] |
| Mind *in vitro* (MiV) | High density | *in vitro (rat – hippocampal; mouse – mESC-derived MNs)* | Zhang (2024)[20] |

**Table 2. User-Specified Parameters in SAMS**

| Parameter | Default Value | Suggested Range * | Description | Refs |
|---|---|---|---|---|
| *Spike Sorting Parameters* | | | | |
| Refractory Period (ms) | 1.5 | 0.5 – 2 | Minimum time elapsing between two consecutive spikes to consider them as coming from the same neuron/unit | 21-24 |
| Threshold_to_Merge | 1.5 | | DTW Distance threshold for an electrode to be flagged as “Possible Multi-Unit” (See **Fig. S1**) | |
| Flag Threshold | 0.1 | | Threshold for the proportion of excluded waveforms during the invalid waveform exclusion procedure for an electrode to be flagged as “Over-Excluded Unit” | |
| Cutoff Frequency (Hz) | 0.1 | 0.01 – 0.2 | Mean firing rate (MFR) cutoff to exclude low-activity neurons/units | 24-27 |
| Standard Deviation Cutoff | 3 | | Number of standard deviations a waveform must exceed from each unit’s mean template in ordered to be considered an outlier and therefore excluded from analysis | |
| *Burst Parameters* | | | | |
| Max ISI (ms) | 100 | 80 - 300 | Maximum Inter-spike Interval (ISI) allowed within a single burst | 23, 28-31 |
| Min # of Spikes | 5 | 4 – 10 | Minimum number of spikes allowed within a single burst | 22, 26, 29 |
| *Network Bursts* | | | | |
| Min # of Spike | 50 | | Minimum number of spikes required in a network burst | 23, 25, 28-31 |
| Max ISI (ms) | 100 | 10 - 300 | Maximum Inter-spike Interval (ISI) allowed within network bursts | 23, 25, 27- 31 |
| Min Electrodes (%) | 35 | 20-80% | The minimum proportion of total electrodes that a burst must occur on in order to be considered a Network Burst | 21, 23, 25, 26, 29-31 |

* All parameters have a range of 0 to +∞ in that SAMS will run without errors with any positive value. Suggested Range values have been taken from published *in vitro* MEA and HD-MEA analyses of hPSC-derived neurons.

**Table S3. Accuracy of SAMS compared to automatic spike sorting algorithms available in OFS software (Related to Figures 4 and S2).**

| Name | Algorithm | Parameters[a] | Correct[b] | Oversort[b] | Undersort[b] |
|---|---|---|---|---|---|
| StdEM (Default) | Standard EM Scan | Seed Cluster Pattern: Circular<br>Beta: 1<br>Max Iterations: 500 | 21.34% (± 1.48%) | 70.23% (± 1.89%) | 8.83% (± 0.81%) |
| StdEM (Alt1) | Standard E-M Scan | **Seed Cluster Pattern: Linear**<br>Beta: 1<br>Max Iterations: 500 | 21.64% (± 1.58%) | 69.48% (± 2.17%) | 8.88% (± 1.04%) |
| StdEM (Alt2) | Standard EM Scan | Seed Cluster Pattern: Circular<br>Beta: 1<br>**Max Iterations: 1000** | **22.09% (± 1.73%)** | 69.18% (± 2.34%) | 8.37% (± 0.96%) |
| TDistEM (Default) | T-Dist E-M Scan | Seed Cluster Pattern: Circular<br>Range of D.O.F. Multiplier: 10 to 30<br>Step by: 5<br>Max. Iterations: 5000<br>Number of Seed Clusters: 5<br>X Remove Outliers | 15.23% (± 6.5%) | 83.86% (± 7.19%) | 0.92% (± 0.7%) |
| TDistEM (Alt1) | T-Dist E-M Scan | Seed Cluster Pattern: Circular<br>**Range of D.O.F. Multiplier: 5 to 25**<br>Step by: 5<br>Max. Iterations: 5000<br>Number of Seed Clusters: 5<br>X Remove Outliers | 15.23% (± 6.5) | 83.86% (± 7.19%) | 0.92% (± 0.7%) |
| TDistEM (Alt2) | T-Dist E-M Scan | Seed Cluster Pattern: Circular<br>Range of D.O.F. Multiplier: 10 to 30<br>Step by: 5<br>Max. Iterations: 5000<br>**Number of Seed Clusters: 4**<br>X Remove Outliers | 4.67% (± 0.09%) | 94.27% (± 0.42%) | 1.06% (± 0.41%) |
| TDistEM (Alt3) | T-Dist E-M Scan | Seed Cluster Pattern: Circular<br>Range of D.O.F. Multiplier: 10 to 30<br>Step by: 5<br>Max. Iterations: 5000<br>Number of Seed Clusters: 5<br>✓ **Remove Outliers** | **41.81% (± 21.5%)** | 56.54% (± 21.87%) | 1.65% (± 0.61%) |
| ValleySeek (Default) | Valley Seeking Scan | Seed Cluster Pattern: Circular<br>Range of Parzen Multiplier: 0.5 to 1.5<br>Step by: 0.2<br>Limit Number of WFs Used to10000<br>✓ Assign unsorted to closest unit<br>✓ Remove Outliers | 19.62% (± 4.04%) | 70.92% (± 5.48%) | 9.45% (± 1.44%) |
| ValleySeek (Alt1) | Valley Seeking Scan | Seed Cluster Pattern: Circular<br>**Range of Parzen Multiplier: 0.3 to 1.3**<br>Step by: 0.2<br>Limit Number of WFs Used to10000<br>✓ Assign unsorted to closest unit<br>✓ Remove Outliers | 9.58% (± 2.57%) | 83.38% (± 4.16%) | 7.04% (± 1.60%) |
| ValleySeek (Alt2) | Valley Seeking Scan | Seed Cluster Pattern: Circular<br>**Range of Parzen Multiplier: 0.7 to 1.7**<br>Step by: 0.2<br>Limit Number of WFs Used to10000<br>✓ Assign unsorted to closest unit<br>✓ Remove Outliers | 14.47% (± 6.63%) | 82.04% (± 7.16%) | 3.49% (± 2.44%) |

| | | | | | |
|---|---|---|---|---|---|
| ValleySeek (Alt3) | Valley Seeking Scan | **Seed Cluster Pattern: Linear**<br>Range of Parzen Multiplier: 0.5 to 1.5<br>Step by: 0.2<br>Limit Number of WFs Used to10000<br>✔ Assign unsorted to closest unit<br>✔ Remove Outliers | **24.54% (± 2.28%)** | 67.09% (± 2.74%) | 8.37% (± 3.05%) |
| K-Means (Default) | K-Means Scan | Seed Cluster Pattern: Circular<br>Range of Units: 2 to 7 | 1.37% (± 0.54%) | 96.68% (± 0.69%) | 1.95% (± 0.53%) |
| K-Means (Alt1) | K-Means Scan | Seed Cluster Pattern: Circular<br>**Range of Units: 1 to 6** | **3.49% (± 1.1%)** | 93.96% (± 1.14%) | 2.55% (± 0.63%) |
| K-Means (Alt2) | K-Means Scan | **Seed Cluster Pattern: Linear**<br>**Range of Units: 1 to 5** | 3.04% (± 0.95%) | 92.91% (± 0.63%) | 4.06% (± 0.4%) |
| SAMS (Pre-Flagged) | SAMS (no user correction of flagged electrodes) | Refractory Period: 1.5<br>Threshold to Merge: 1.5<br>Flag Threshold: 0.1<br>Cutoff Frequency: 0.1<br>Standard Deviation Cutoff: 3 | 84.32% (± 2.34%) | 10.54% (± 1.82%) | 5.13% (± 0.6%) |
| SAMS (Post-Flagged) | SAMS (with user correction of flagged electrodes) | Refractory Period: 1.5<br>Threshold to Merge: 1.5<br>Flag Threshold: 0.1<br>Cutoff Frequency: 0.1<br>Standard Deviation Cutoff: 3 | **92.64% (± 2.1%)** | 4.21% (±1.24%) | 3.16% (± 0.91%) |

[a] Bold font indicates parameter that was changed from the default value
[b] Values shown are mean percent of total electrodes (± SEM) for n =3 data files (same files used for analyses shown in **Figure 4**).

**Table S4. Comparison of MEA Parameters in Data from Shen, Sirois, Guo, Li et al (2023)[31] Before and After Spike-Sorting Using SAMS**

| Parameter | Standard MEA Analysis | Results After Spike-Sorting |
|---|---|---|
| Mean Firing Rate | **↑ in FXS2 **** | **↑ in FXS2 ****** |
| Burst Duration | **n.d.** | **↓ in FXS2 ****** |
| Number of Spikes per Burst | **↑ in FXS2 *** | **↑ in FXS2 ****** |
| Mean ISI in Burst | **n.d.** | **↓ in FXS2 ****** |
| Inter-burst Interval (IBI) | **n.d.** | **↓ in FXS2 ****** |
| Burst Frequency | **↑ in FXS2 *** | **↑ in FXS2 ****** |
| Normalized Duration IQR | **n.d.** | **↑ in FXS2 ****** |
| IBI Coefficient of Variation (CoV) | **n.d.** | **n.d.** |
| Burst Percentage | **↑ in FXS2 *** | **↑ in FXS2 ****** |
| Network Burst Frequency | **↑ in FXS2 **** | **↑ in FXS2 ***** |
| Network Burst Duration | **n.d.** | **↓ in FXS2 *** |
| Number of Spikes per Network Burst | **↑ in FXS2 **** | **↑ in FXS2 ****** |
| Network Burst Percentage | **↑ in FXS2 *** | **↑ in FXS2 ****** |
| Network Burst IBI CoV | **n.d.** | **n.d.** |

Comparison of FXS2 versus CTRL2 at the 7 week time point. Unpaired t-test (no significant difference in variance between groups) or Welch's t-test (significant difference in variance between groups) * $p < 0.05$, ** $p < 0.01$, *** $p < 0.005$, **** $p < 0.001$ Standard MEA analysis: N = 4 independent differentiations; Results After Spike-Sorting: n = 298 neurons (CTRL2) and 837 neurons (FXS2) identified through spike sorting from N = 4 independent differentiations, with n = 85 CTRL2 and 685 FXS2 neurons exhibiting bursting activity, and n = 11 CTRL2 and 49 FXS2 neurons exhibiting network bursting. After removal of outliers, n = 249-759 (Mean Firing Rate), n = 75-627 (bursting parameters), n = 11-48 (network bursting parameters)

### Table S5. Electrode and Well Information for All Data Files Analyzed

<table>
<tr><th rowspan="2">File Name</th><th rowspan="2">Figure(s)</th><th rowspan="2">Experimental Conditions in File</th><th colspan="2">Wells</th><th colspan="2">Electrodes</th></tr>
<tr><th>Recorded</th><th>Active</th><th>Recorded</th><th>Active</th></tr>
<tr><td>File 1[c]</td><td>4, 5, 6, 7, S2, S3, S4, Tables S3 & S4</td><td>FXS2 and CTRL2 neurons</td><td>48</td><td>46</td><td>384</td><td>220</td></tr>
<tr><td>File 2[c]</td><td>4, 5, 6, 7, S2, S3, S4, Tables S3 & S4</td><td>FXS2 and CTRL2 neurons</td><td>96</td><td>69</td><td>768</td><td>219</td></tr>
<tr><td rowspan="2">File 3[b, c]</td><td>4, 5, S2, S3, Table S3</td><td>FXS2 and CTRL2 neurons</td><td>48</td><td>43</td><td>384</td><td>225</td></tr>
<tr><td>6, 7, S4, Table S4</td><td>FXS2 and CTRL2 neurons</td><td>48</td><td>48</td><td>384</td><td>288</td></tr>
<tr><td>File 4[c]</td><td>6, 7, S4, Table S4</td><td>FXS2 and CTRL2 neurons</td><td>36</td><td>29</td><td>288</td><td>97</td></tr>
<tr><td>File 5[d]</td><td>6, 7</td><td>CTRL1/shNC, FXS1/shNC, and FXS1/shMAP1B neurons [NGN2-differentiation]</td><td>72</td><td>67</td><td>576</td><td>300</td></tr>
<tr><td>File 6[d]</td><td>S4</td><td>CTRL3 (dCas9-activator + scramble sgRNA) and MAP1B-EE (dCa9-activator + MAP1B sgRNA) neurons [NGN2-differentiation]</td><td>96</td><td>96</td><td>768</td><td>602</td></tr>
</table>

**a** Active refers to wells or electrodes that contain active neurons after running SAMS pipeline (units with a minimum mean firing rate > 0.1 Hz)

**b** The top half of this row refers to the wells and electrodes used to assess the accuracy of SAMS (Figures 4, 5, S2, S3), while the bottom half of this row refers to wells and electrodes used to compare activity in FXS2 and CTRL2 neurons (Figures 6, 7, S4).

**c** File from Shen, Sirois, Guo, Li et al (2023)[31]

**d** File from Guo et al (2023)[30]

**Table S6. Settings Used for Spike File Generation (AxIS Navigator Software)**

| | |
|---|---|
| **Method** | |
| Threshold: | 6 x Standard Deviation |
| Detect Only Crossings: | Yes |
| Peak Detection: | Maximum Amplitude |
| **Durations** | |
| Pre-Spike: | 0.84 ms (11 samples) |
| Post-Spike: | 2.16 ms (27 samples) |
| Allow Sample Overlap: | No |
| **Coincident Events** | |
| Coincident Event Removal: | Well |
| **Spike Counting** | |
| Interval: | 1s (12.5 K samples) |
